# T cell derived HB-EGF prevents Th17 cell differentiation in an autocrine way

**DOI:** 10.1101/2021.02.09.430418

**Authors:** Felicity Macdonald, Jorg van Loosdregt, Dietmar M W Zaiss

## Abstract

CD4 T cells critically contribute to host immunity against infections, but can also contribute to the development of autoimmune diseases. The underlying mechanisms that govern differentiation of naïve CD4 T cells into different effector populations remain poorly understood. Here, we show that the expression of the Epidermal Growth Factor (EGF)-like growth factor HB-EGF by CD4 T cells sustained their expression of Interleukin (IL)-2 and reduced their capacity to differentiate into T Helper 17 (Th17) cells. Concordantly, mice with a T cell specific deficiency of HB-EGF showed an enhanced differentiation of naïve CD4 T cells into Th17 cells and a more rapid onset of experimental autoimmune encephalomyelitis (EAE). Furthermore, transfer of naïve HB-EGF-deficient CD4 T cells into *Rag1^-/-^* mice led to the rapid induction of multi-organ inflammation in recipient mice. Together, our data reveal a novel mechanism by which an HB-EGF-mediated constrain on Th17 differentiation prevents the development of autoimmune diseases.

**SUMMARY:** CD4 T cell activation induces the expression of the EGFR and its high-affinity ligand HB-EGF. HB-EGF sustains IL-2 expression in an autocrine manner, preventing the differentiation of Th17 cells and the subsequent induction of Th17 cell-mediated autoimmune diseases.

## INTRODUCTION

CD4 T cells are key regulators of immune responses. Cytokines produced by CD4 T cells during pathogen-specific responses are critical effector molecules that regulate the local immune response and pathogen clearance. For instance, IFNγ-producing Th1 cells directly contribute to the clearance of *Leishmania* or *Mycobacterium leprae* (Modlin, 1994), IL-17-producing Th17 cells contribute to fungal resistance (Conti et al., 2009), and IL-13-producing Th2 cells contribute to the clearance of helminths (Else et al., 1994; Minutti et al., 2017). Deficiencies in T cell differentiation can result in enhanced pathogen load and immune pathology. In addition, T cells can contribute to the development of autoimmune diseases and the differentiation of naïve CD4 T cells into pro-inflammatory Th17 cells has been shown to be a critical step in the development of autoimmune diseases (Annunziato et al., 2009; Ivanov et al., 2006). Therefore, the correct differentiation of CD4 effector populations is a fundamental step in the regulation of immune responses. Nevertheless, the precise mechanisms that direct *in vivo* CD4 T cell differentiation has remained poorly understood.

We and others have shown that Epidermal Growth Factor Receptor (EGFR) expression is essential for efficient regulatory T cell (Treg) and effector CD4 T cell function and therefore critically contributes to immune regulation (Zaiss et al., 2013), host defense (Minutti et al., 2017) and the development of immune mediated diseases, such as atherosclerosis (Zeboudj et al., 2018). Nevertheless, whether EGFR expression by CD4 T cells may also influence their differentiation into different effector types, has remained unresolved so far.

The EGFR is an evolutionary conserved transmembrane receptor known to play a critical role in tissue development and cancer formation (Sibilia et al., 2007). The EGFR is also widely expressed by leukocytes, such as macrophages (Hardbower et al., 2016; Lanaya et al., 2014) and plasma cells (Mahtouk et al., 2005), and is a critical regulator of stress-induced expression of NKG2D ligands such as MICA/B, which govern the local initiation of immune responses by activating NK-cells (Vantourout et al., 2014). Furthermore, human and mouse CD4 T cells express the EGFR upon activation (Minutti et al., 2017; Zaiss et al., 2013; Zeboudj et al., 2018). The EGFR has been shown to be one of the most upregulated genes upon STAT-5 signalling in mouse CD4 T cells (Beier et al., 2012; Liao et al., 2008) and, in human Tregs, the EGFR was found to be the single most upregulated transmembrane kinase upon activation (Tuettenberg et al., 2016).

Similar to the EGFR, the high-affinity EGFR ligand Heparin-Binding EGF-like Growth Factor (HB-EGF) has been shown to be expressed on newly-activated naïve CD4 T cells (Blotnick et al., 1994). HB-EGF binding leads to the autophosphorylation of the EGFR, ensuing MAP-kinase (MAPK) and STAT-5 signalling pathway activation (Heo et al., 2018). Both signal transduction pathways are crucial for the differentiation of CD4 T cells. In addition, upon endocytosis induced by the binding of high-affinity ligands, EGFR activity is closely regulated by the phosphate PTPN-2 (Stanoev et al., 2018). Since individuals with dysfunctional PTPN-2 expression are strongly predisposed to the development of auto-immune diseases, and as this phosphate is highly expressed in T cells (Doody et al., 2009), we decided to test whether EGFR mediated signalling and the expression of its ligands by CD4 T cells may influence the differentiation of CD4 T cells into their different subpopulations and in this way may contribute to the development of autoimmune diseases.

Here, we describe the finding that the high-affinity EGFR ligand HB-EGF blocks Th17 differentiation in an autocrine manner and that both *in vitro* and *in vivo* HB-EGF-deficient CD4 T cells differentiated more readily into Th17 cells. Upon transfer into *Rag1^-/-^* mice, recipients of naïve HB-EGF-deficient CD4 T cells developed unanticipated early-onset of wasting disease and multi-organ inflammation, which also affected the central nervous system. Mechanistically, we found that CD4 T cells deficient in HB-EGF express lower levels of IL-2 upon activation and showed diminished STAT-5 signalling. As STAT-5 signalling has been shown to interfere with Th17 differentiation (Laurence et al., 2007), these findings suggest a novel mechanism by which HB-EGF-mediated signalling in naïve CD4 T cells forms a threshold preventing their aberrant differentiation into Th17 cells, thereby limiting the development of autoimmune diseases.

## RESULTS

### Unaltered T cell compartment in mice with T cell-specific HB-EGF deficiency at steady state

HB-EGF has been described to be expressed by CD4 T cells following activation (Blotnick et al., 1994). To confirm these findings in CD4 T cells derived from C57BL/6 mice, we activated naïve CD4 T cells either using α-CD3/α-CD28 coated beads or naïve CD4 T cells from from a TCR transgenic mouse line in peptide-specific way. We detected a dose-dependent expression of HB-EGF on activated CD4 T cells (Figure 1A & B). We have shown before that EGFR expression by CD4 T cells can be sustained by STAT-5 inducing cytokines, but initially requires an induction of expression via the TCR (Minutti et al., 2017). Remarkably, in the case of HB-EGF deficient CD4 T-cells we found that induced HB-EGF expression was essential for the induction of EGFR expression (Figure 1C). Based on these data, we concluded that an HB-EGF mediated feed-back loop is essential for the induction of appreciable levels of EGFR expression on newly activated CD4 T cells.

**Figure 1:**
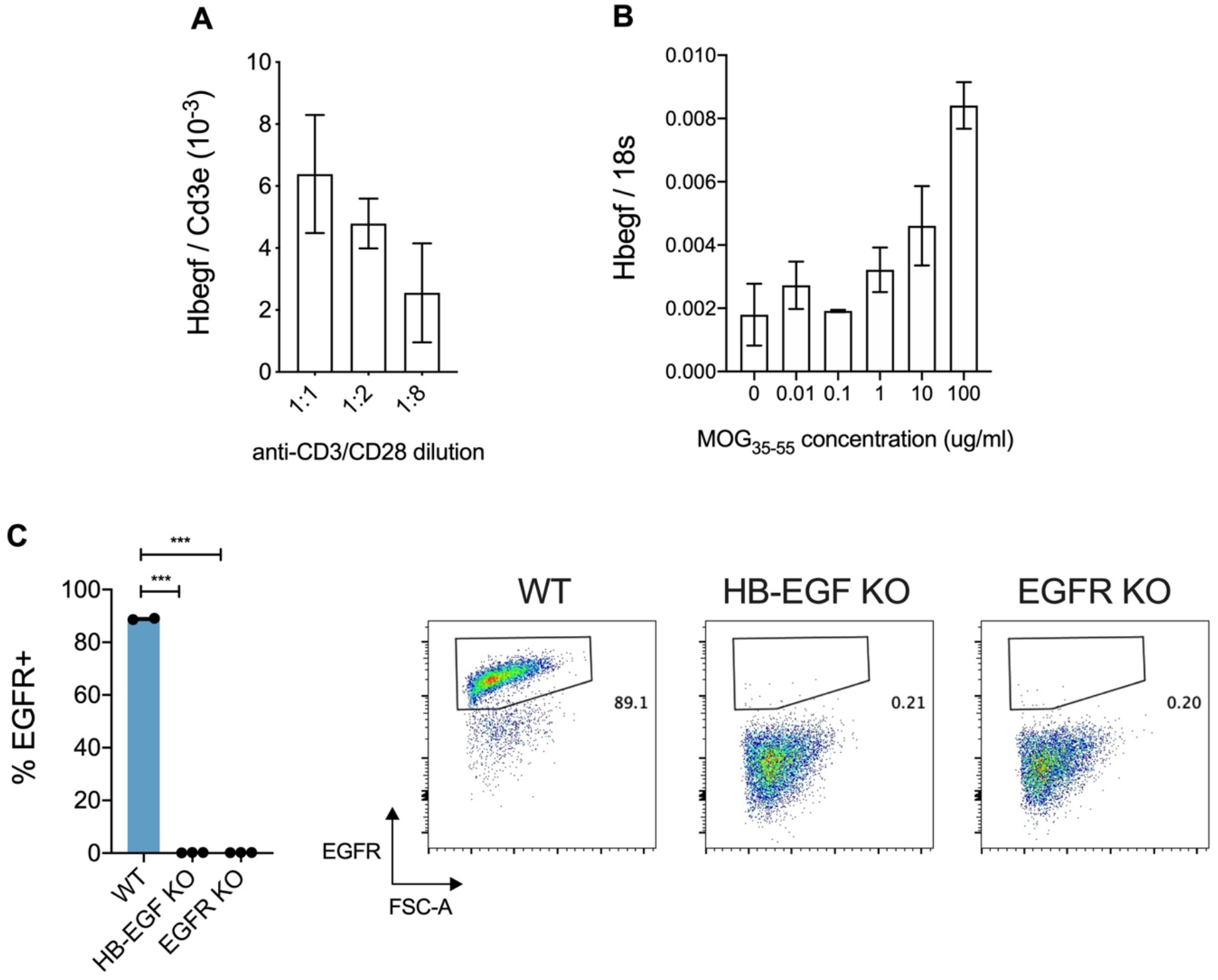
CD4+ T cells express EGFR and HB-EGF upon activation. (A) Naïve CD4+ T cells were activated with decreasing concentrations of α-CD3/α-CD28 stimulatory beads *in vitro*. Quantification of *Hbegf* mRNA expression was performed following o/n stimulation. (B) Naïve 2D2 TCR transgenic CD4+ T cells were activated *in vitro* with BM-DCs pulsed with increasing concentrations of TCR specific MOG peptide. Quantification of *Hbegf* mRNA expression is shown following o/n stimulation. (C) Naïve CD4+ T cells from WT, CD4D^HB-EGF^ and CD4^DEGFR^ mice were activated *in vitro* with α-CD3/α-CD28-coated stimulator beads for 24 hours. Cells were stained for CD4 and EGFR, and analysed by flow cytometry. Surface staining and graphs show quantification of EGFR expression. Error bars show mean ± SEM. n=2-3 in each group.

In order to evaluate the function of T cell-expressed HB-EGF, we established mice with a T cell-specific deficiency in HB-EGF (*CD4:cre x Hbegf^fl/fl^*, from now on called CD4^ΔHB-EGF^). To ascertain the activation states of T cells in these mice at steady state, we analysed T cells derived from spleen, inguinal lymph nodes (iLN), intraepithelial lymphocytes (IEL) and the lamina propria (LP) of the small intestine for CD44 and CD62L expression. As shown in Figure S1, we saw no difference in number and percentage of naïve CD4^+^ T cells (CD62L^+^ CD44^-^), effector CD4^+^ T cells (CD62L^-^ CD44^+^) or central memory CD4^+^ T cells (CD62L^+^ CD44^+^) compared to wild type (wt) C57BL/6 mice and mice with a T cell-specific deficiency in EGFR expression (*Cd4:cre x Egfr^fl/fl^*, from now on called CD4^ΔEGFR^).

These data suggest that neither a T cell specific lack of HB-EGF or of EGFR expression influences the thymic maturation of T cell subpopulations or induce the development of spontaneous autoimmune diseases at steady state. Therefore, we concluded that similar to our finding in CD4^ΔEGFR^ mice (Minutti et al., 2017), T cell-specific HB-EGF deficiency does not play a major role in T cell development and homeostasis.

### Rapidly expansion of Th17 cell populations in CD4^ΔHB-EGF^ mice

To assess the influence of T cell-derived HB-EGF on *in vivo* CD4 T cell differentiation, we injected activating αCD3 antibodies into wt C57BL/6 and CD4^ΔHB-EGF^ mice. It has previously been reported that αCD3 antibody injection induces activation-induced T cell death, along with a systemic upregulation of IL-6 and TGFβ. This induces the *de novo* differentiation of Th17 cells, their recruitment to the small intestine and local inflammation in the small intestine (Esplugues et al., 2011). As shown in Figures 2, we found that 48 hours following injection of αCD3 antibodies into wt C57BL/6 and CD4^ΔHB-EGF^ mice the proportion of IL-17^+^, RORγt^+^ and IL-17^+^ RORγt^+^ double-positive IELs in CD4^ΔHB-EGF^ mice was significantly increased compared to wt counterparts.

**Figure 2:**
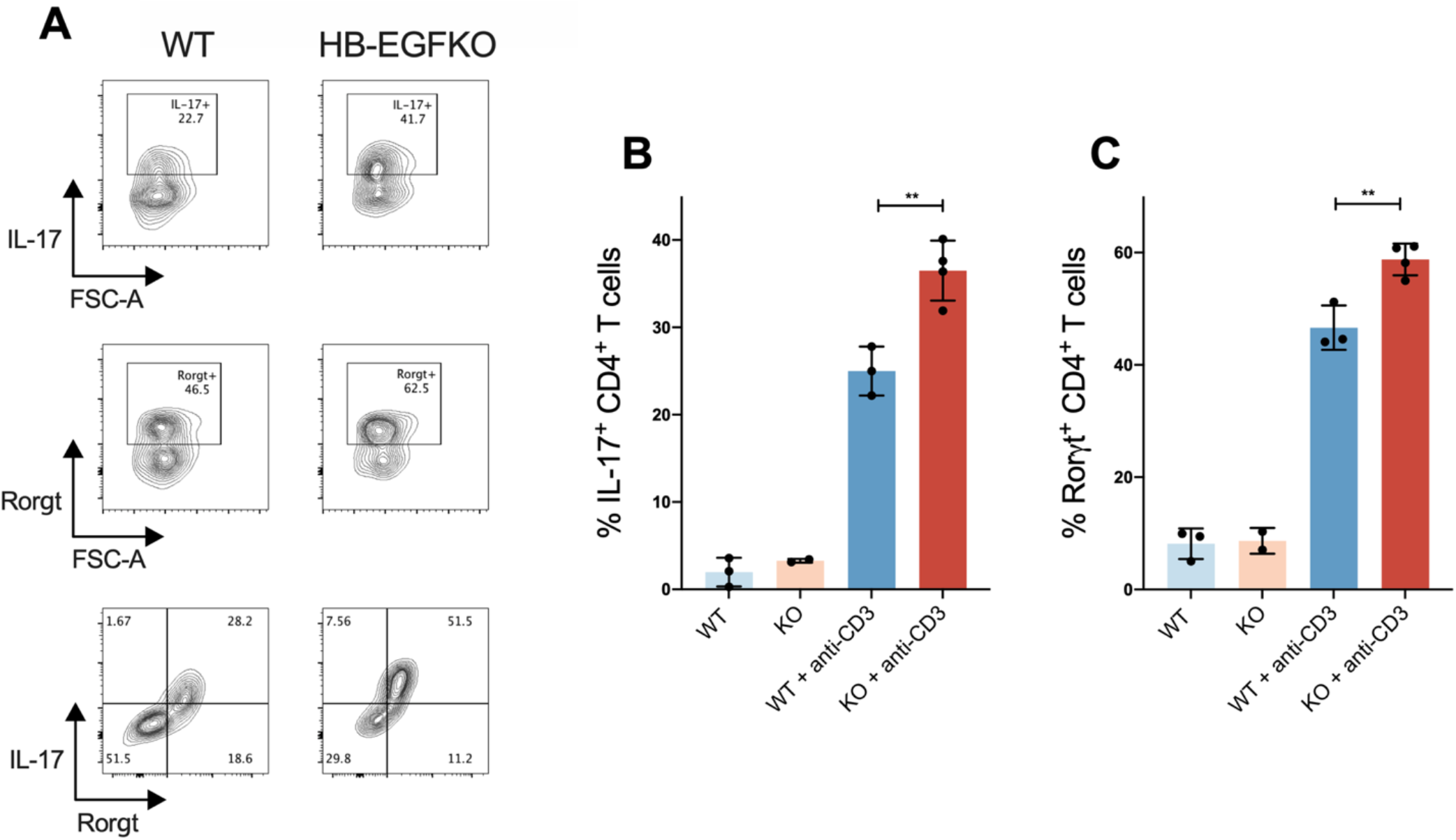
Enhanced *in vivo* expansion of the Th17 population in mice with HB-EGF-deficient T cells using a model of α-CD3-induced T cell activation. (A-C) WT and CD4^ΔHB-EGF^ mice were injected intraperitoneally with α–CD3 antibody, and culled 48 hours later. Intraepithelial lymphocytes (IEL) were isolated from small intestines. Cells were stained for CD4, CD3, IL-17 and Rorγt before analysis by flow cytometry. (A) Representative FACS plots and quantified data is shown for IL-17^+^ cells (top), Rorγt^+^ T cells (middle) and IL-17^+^ Rorγt^+^ T cells (bottom) as a proportion of CD4^+^ CD3^+^ lymphocytes. (B+C) Quantified data seen in (A) for IL-17^+^ cells (B) or Rorγt^+^ T cells (C) as a proportion of CD4^+^ CD3^+^ lymphocytes. Error bars show mean ± SEM. n=2-5 in each group. Student’s t tests were used to assess significance.

These data suggest that a deficiency of HB-EGF expression by CD4 T cells enhances the *in vivo* differentiation of Th17 cells.

### HB-EGF suppresses Th17 cell differentiation *in vitro*

To determine whether HB-EGF-deficient CD4 T cells retain the same capacity as wt cells to differentiate into different effector populations, we sorted naïve CD4 T cells from C57BL/6 wt, CD4^ΔEGFR^ and CD4^ΔHB-EGF^ mice and polarised these cells *in vitro*. As shown in Figure 3, we found that similar to Th2 cells (Minutti et al., 2017) also Th1 and induced Treg (iTreg) differentiation does not appear to be substantially influenced by a lack of HB-EGF or EGFR expression. However, we detected enhanced Th17 cell differentiation with both EGFR and HB-EGF deficiency when staining for the Th17 master transcription factor RORγt (Figure 3A). In the presence of recombinant IL-6 and TGFβ, a significantly higher percentage of CD4 T cells derived from CD4^ΔEGFR^ and CD4^ΔHB-EGF^ mice differentiated into Rorγt-expressing Th17 cells compared to CD4 T cells derived from wt C57BL/6 mice (Figure 3A). This increased potential to differentiate into Th17 cells was reversed in a dose-dependent manner upon the addition of recombinant HB-EGF to the polarisation culture media of CD4 T cells derived from CD4^ΔHB-EGF^ mice, but not in those derived from CD4^ΔEGFR^ mice (Figure 3D). These data suggest that HB-EGF constrains the differentiation of naïve CD4 T cells into Th17 cells by signalling through the EGFR.

**Figure 3:**
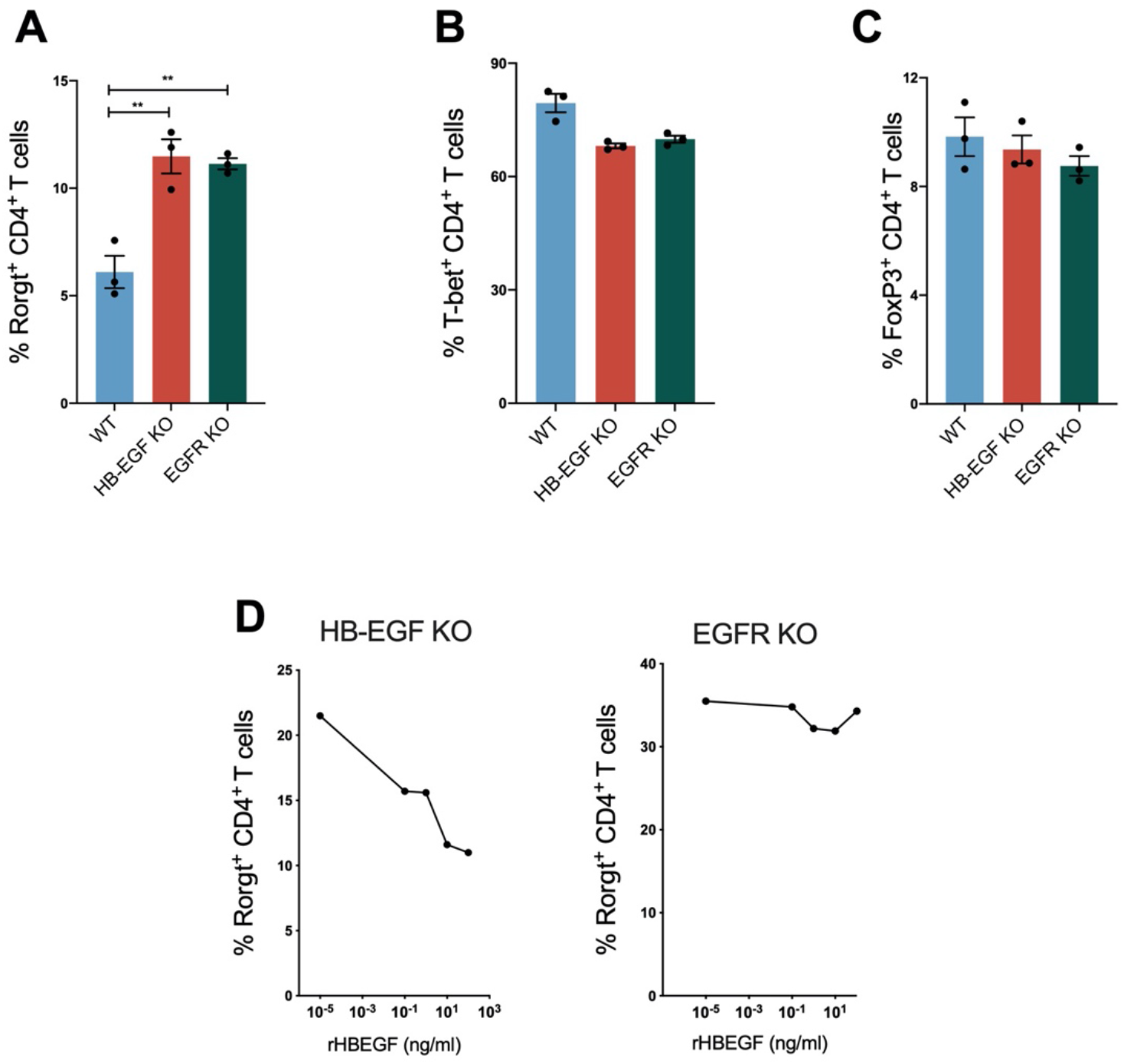
Enhanced *in vitro* Th17 cell differentiation of HB-EGF-deficient CD4+ T cells. Naïve CD4^+^ T cells (CD4^+^ CD45RB^+^ CD25^-^) were FACS sorted from spleens of WT, CD4^ΔHB-EGF^ and CD4^ΔEGFR^ mice and activated *in vitro* using α-CD3/α-CD28 antibodies coated beads. (A) Cells were cultured in IL-1 β, IL-6, IL-23, TGFβ and FICZ for 5 days to polarise towards a Th17 phenotype. Cells were stained for CD4, CD3 and Th17 master transcription factor RORγt before analysis by flow cytometry. Dead cells were excluded from analysis by staining cells with a live/dead dye. Quantification of Rorγt^+^ cells as a proportion of CD4^+^ CD3^+^ lymphocytes. (B+C) Cells were polarised towards a Th1 (IL-12 and α-IL-4) (B) or iTreg (TGFβ and IL-2) (C) phenotype. Cells were stained for CD4, CD3 and master transcription factors T-bet (Th1) or FoxP3 (iTreg) before analysis by flow cytometry. Graphs show quantification of T-bet^+^ (B) or FoxP3^+^ (C) cells as a proportion of CD4^+^ CD3^+^ lymphocytes. (D) Recombinant HB-EGF was added to the culture media of naïve CD4+ T cells before polarisation towards a Th17 phenotype as in (A). Graphs show quantification of Rorγt^+^ cells as a proportion of CD4^+^ CD3^+^ lymphocytes upon addition of increasing concentrations of rHB-EGF from CD4^ΔHB-EGF^ (left) or CD4^ΔEGFR^ (right) mice. Error bars show mean ± SEM. n=3 in each group. Student’s t tests were used to assess significance.

To assess whether human CD4 T cells respond to HB-EGF in a similar fashion, PBMCs were obtained from healthy donor blood and CD4 T cells were isolated. Cells were bulk transfected with recombinant CRISPR/Cas9 and guide RNA specific for *Hbegf* or *Egfr* gene sequences or a scrambled control guide RNA. After resting for two days, cells were *in vitro* polarised towards a Th17 phenotype. While we only found a partial reduction in HB-EGF expression in CRISPR/Cas9 treated cells by Western blot of bulk treated cells, we consistently found a significantly higher percentage of Rorγt^+^ Th17 cells in CD4 T cell populations treated with *Hbegf* or *Egfr* gene-specific guide RNA than in unstimulated CD4 T cells or CD4 T cells which underwent editing using a scrambled guide RNA (Figures S2A & B). These findings suggest that *in vitro* HB-EGF-mediated signalling has a similar ability to diminish Th17 cell differentiation in both mouse and human CD4 T cells.

### HB-EGF enhances IL-2 expression in activated CD4 T cells

In order to better understand the mechanism by which HB-EGF-mediated signalling diminished Th17 cell differentiation, we measured the production of IL-2 by activated CD4 T cells. We have shown previously that EGFR-deficient CD4 T cells initially show inferior proliferation upon activation compared to their wt counterparts (Minutti et al., 2017). Activated CD4 T cells require autocrine IL-2 to aid in clonal expansion, and the proliferation deficiency of EGFR-deficient CD4 T cells could be overcome by the supplementation of culture media with recombinant IL-2 in a dose dependent manner (Minutti et al., 2017).

To investigate the influence of T cell derived HB-EGF on IL-2 expression, we activated naïve CD4 T cells *in vitro* and measured proliferation along with IL-2 production over the following 24-48 hours. We found that in the presence of recombinant IL-2, proliferation of T cells derived from CD4^ΔHB-EGF^ or C57BL/6 wt mice was comparable (Figure 4A). However, both HB-EGF- and EGFR-deficient CD4 T cells produced significantly less IL-2 than wt cells upon activation (Figure 4B). In line with these findings, the activation-induced upregulation of the IL-2 receptor, CD25, was significantly diminished, as was the activation of the STAT-5 signalling pathways, in HB-EGF- and EGFR-deficient CD4 T cells (Figure 4C & D).

**Figure 4:**
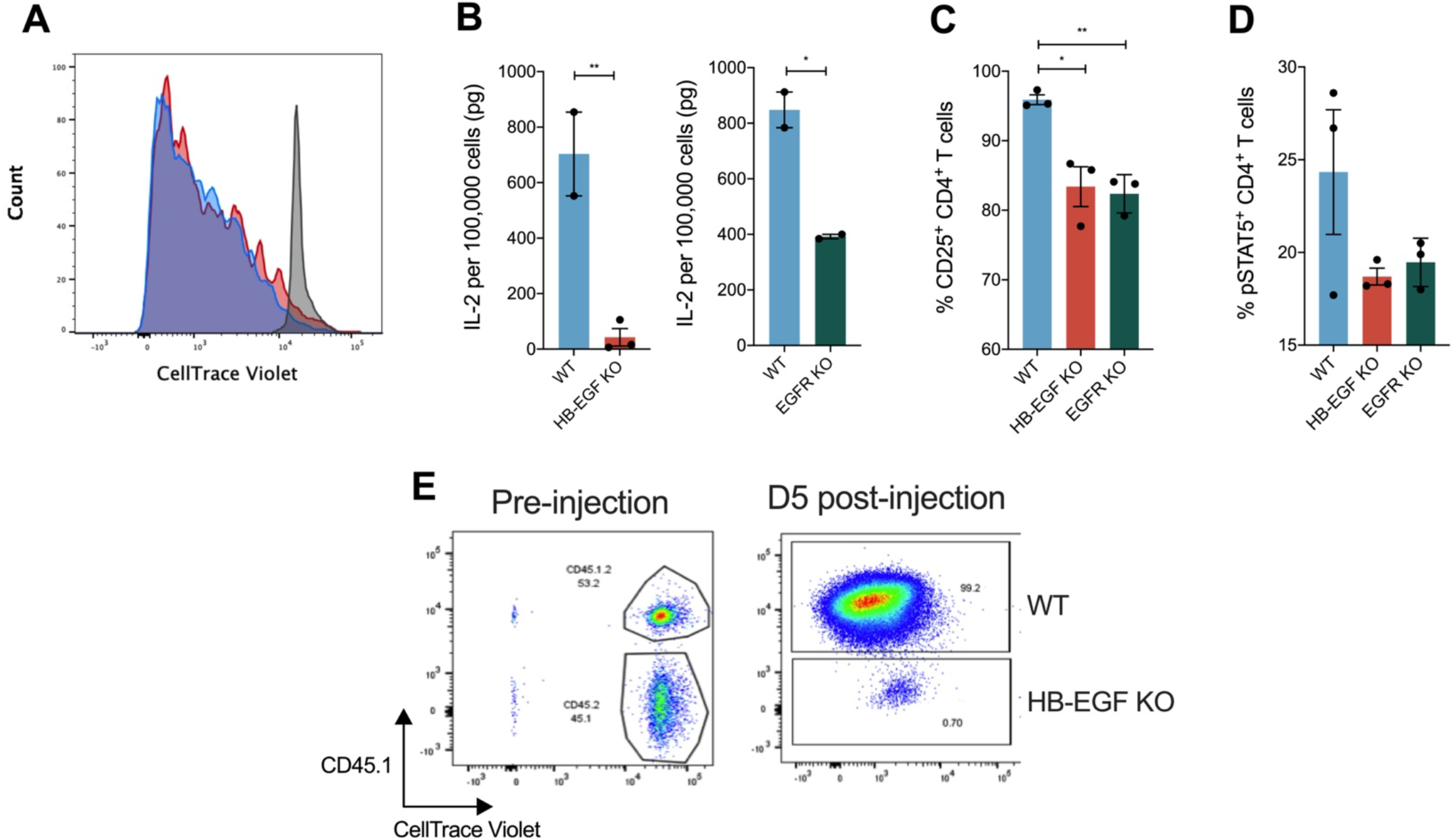
HB-EGF-deficient CD4 T cells produce less IL-2 upon activation and show inferior recovery compared to wt cells after transfer into *Rag1^-/-^* mice. (A) Sorted naïve CD4^+^ T cells from spleens of wt and CD4^ΔHB-EGF^ mice were stained with CellTrace Violet Cell Proliferation Kit and activated *in vitro* in the presence of recombinant IL-2 for 4 days. FACS plot shows unstimulated control cells after 24 hours in culture (grey), along with cells from WT (blue) and CD4^ΔHB-EGF^ (red) mice. Each peak represents distinct proliferation. (B-D) Naïve CD4^+^ T cells from WT, CD4^ΔHB-EGF^, and CD4^ΔEGFR^ mice were activated *in vitro* with α-CD3/α-CD28-coated stimulator beads for 24-48 hours. (B) Supernatants were harvested, and ELISA was performed to determine IL-2 concentration per 100,000 cells. (C+D) Cells were stained for CD4, CD3, CD25 and intranuclearly for pSTAT5, and analysed by flow cytometry. Surface staining and graphs show quantification of CD25^+^ (C) or pSTAT5^+^ (D) cells as a proportion of CD4^+^ CD3^+^ lymphocytes. (E) *Rag1^-/-^* mice (CD45.1) were intravenously injected with a 1:1 mixture of WT (CD45.1/CD45.2) and HB-EGF-deficient (CD45.2) naïve CD4 T cells. FACS plot of WT (CD45.1^+^) and HB-EGF-deficient (CD45.1^-^) cell mixture post injection (left) or 5 days after injection into *Rag1^-/-^* mice against CellTrace Violet staining (right). All experiments were performed at least twice with similar results. Error bars show mean ± SEM. n=2-3 in each group. Student’s t tests were used to assess significance.

To further explore the proliferative capacity of HB-EGF-deficient T cells *in vivo*, we utilised the *Rag1^-/-^* transfer model, in which intravenously injected naïve CD4 T cells undergo lymphopenia-induced proliferation and activation (Bourgeois et al., 2005). Upon injection of a 1:1 mix of wt (CD45.1/CD45.2) and CD4^ΔHB-EGF^ (CD45.2) naïve CD4 T cells into *Rag1^-/-^* mice, we observed a dramatically diminished recovery of HB-EGF-deficient cells in comparison to wt CD4 T cells on day 5 post transfer, with the majority of retrieved donor cells coming from the transferred wt T cell population (Figure 4E).

These data illustrate that HB-EGF-mediated signalling sustains CD4 T cell expansion in an autocrine manner, potentially by enhancing the production of IL-2 by activated CD4 T cells.

### Transfer of naive HB-EGF-deficient CD4 T cells into *Rag1^-/-^* mice induces their rapid differentiation into Th17 cells

IL-2-induced STAT-5 signalling has been shown before to interfere with Th17 cell differentiation (Laurence et al., 2007). Therefore, our finding that HB-EGF-deficient CD4 T cells express diminished levels of IL-2 may account for the observed enhanced *in vitro* differentiation into Th17 cells. To further assess the capacity of naïve HB-EGF-deficient CD4 T cells to differentiate into Th17 cells *in vivo*, we followed the transferred 1:1 mix of wt (CD45.1) and HB-EGF-deficient (CD45.2) T cells in *Rag1^-/-^* mice for an extended period of time. As time progressed, we found that the transferred HB-EGF-deficient T cell population recovered until there was an equal ratio of CD45.1^+^ wt to CD45.2^+^ HB-EGF-deficient donor CD4 T cells (Figure 5A). Furthermore, when the transferred populations were assessed for effector cell differentiation, we found that from day 14 post-transfer onwards there was a significantly stronger polarisation of IL-17 and IFNγ co-secreting Th17 cells within the CD45.2^+^ HB-EGF-deficient T cell population compared to the CD45.1^+^ wt T cell population (Figure 5B), demonstrating that *in vivo* CD4^ΔHB-EGF^ cells differentiated more readily into Th17 cells than their co-transferred wt counterparts.

**Figure 5:**
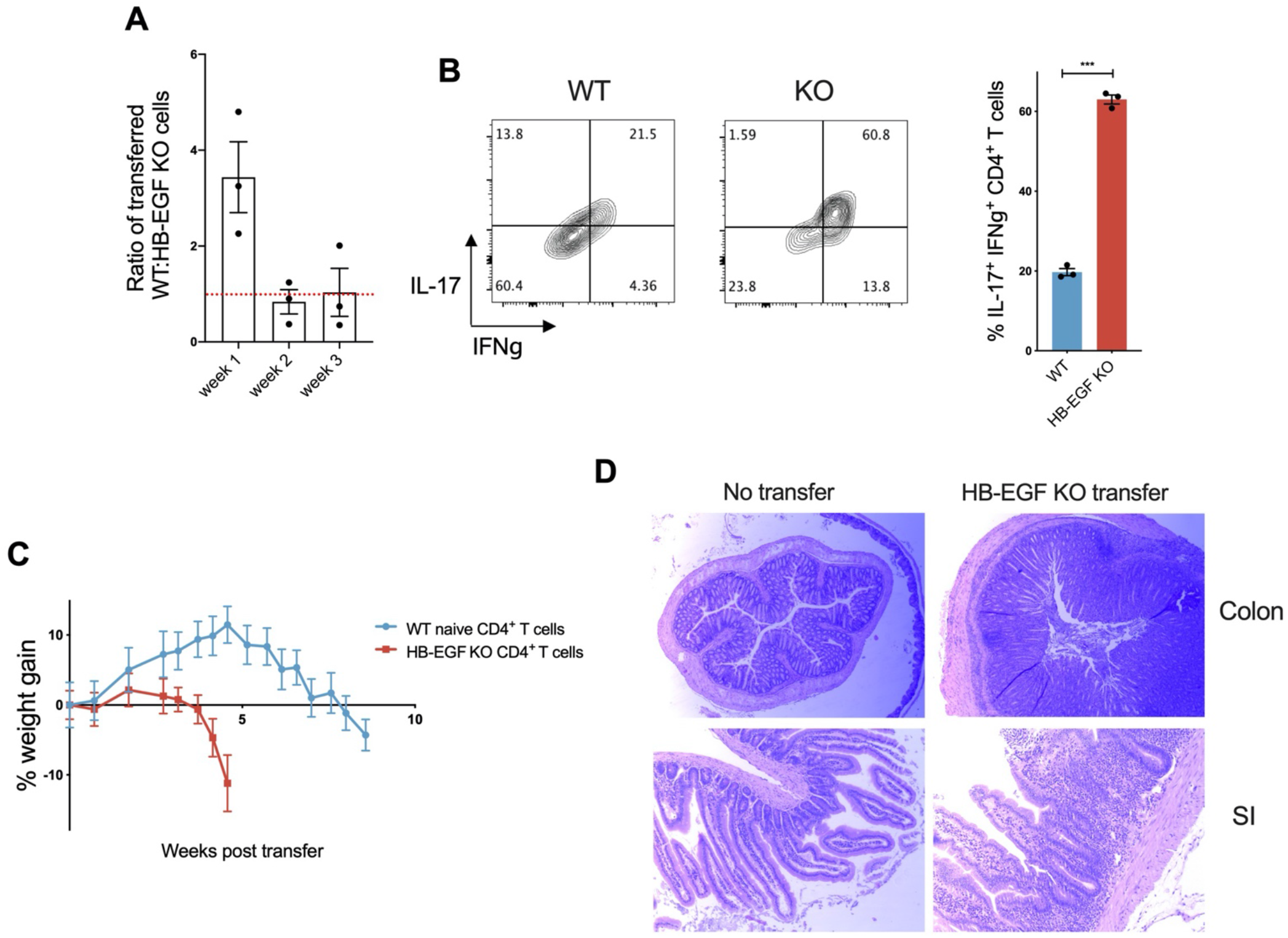
HB-EGF-deficient CD4 T cells show enhanced Th17 differentiation and cause early onset of wasting disease. (A-B) *Rag1^-/-^* mice (CD45.1) were intravenously injected with a 1:1 mixture of WT (CD45.1/.2) and HB-EGF-deficient (CD45.2) naïve CD4 T cells. (A) Ratio of WT (CD45.1) to HB-EGF-deficient (CD45.2) CD4 T cells in recipient *Rag1^-/-^* mice at different time points post transfer. (B) Mice were culled at week 1 post transfer, and splenocytes stimulated *ex vivo* with PMA and ionomycin for 4 hours. Cells were subsequently stained for CD45.1, CD45.2, CD4, CD3 plus intracellularly for IL-17 and IFNγ. FACS plots and quantified data show percentage of IL-17^+^ IFNγ^+^ co-expressing T cells as a proportion of CD4^+^ lymphocytes. (C) *Rag1^-/-^* mice were intravenously injected with either 500,000 WT (blue) or HB-EGF-deficient (red) naïve CD4 T cells. Mice were weighed frequently for onset of wasting disease. Graph shows percentage weight gain of mice compared to their original body weight post injection. (D) Histology slides of colon (x20 magnification) or small intestine (x100) from either C57Bl/6 wt mice or *Rag1^-/-^* mice injected with HB-EGF-deficient naïve CD4 T cells (taken 3 weeks post-transfer) stained with H&E. Error bars show mean ± SEM. n=3-5 in each group. Student’s t tests were used to assess significance.

These data further support the proposition that T cell-derived HB-EGF counteracts the differentiation of naïve CD4 T cells into Th17 cells in an autocrine manner.

### Transfer of naive HB-EGF-deficient CD4 T cells into *Rag1^-/-^* mice induces the rapid onset of multi-organ inflammation and wasting disease

Transfer of naïve CD4 T cells into lymphopenic mice is a well-established model for the induction of colitis (Powrie et al., 1994). For colitis induction, transferred T cells have first to differentiate into Th17 cells before they migrate in a CXCR6-dependent manner into the colon, where they then differentiate into IFNγ^+^ Th1/Th17-like cells and cause colitis (Harbour et al., 2015). The transfer of fully differentiated Th17 cells into lymphopenic mice causes a rapid onset of disease (Lee et al., 2009). Therefore, to assess the capacity of HB-EGF-deficient T cells to induce colitis, we transferred naïve CD4 T cells derived from either C57BL/6 wt or CD4^ΔHB-EGF^ mice into *Rag1^-/-^* mice. The injection of naïve wt CD4 T cells induced the expected progression of disease in recipient *Rag1^-/-^* mice, with weight loss being initiated at around 6 weeks after transfer (Harbour et al., 2015). However, the transfer of naïve HB-EGF-deficient CD4 T cells induced an unexpected, rapid onset of wasting disease at around 3 weeks post transfer (Figure 5C). The disease was characterized by multi-organ inflammation, which was especially pronounced in the small and large intestine (Figure 5D). However, in these mice we occasionally also observed the onset of atypical EAE (constant turning of individuals due to a loss of balance) as well as conventional EAE (full paralysis of tail and hind legs). To test whether TGFβ-mediated differentiation of transferred T cells into Th17 cells might be contributing to the early onset of disease in recipient mice, we injected TGFβ blocking antibodies (1D11) into *Rag1^-/-^* mice that had received naïve CD4 T cells from CD4^ΔHB-EGF^ mice. Injection of TGFβ blocking antibodies prevented the early onset of disease (Figure S3A), suggesting that TGFβ mediated differentiation into Th17 cells contributes to the early onset of disease. As we did not detect a difference in the differentiation of transferred naïve CD4 T cells into iTregs between the mouse groups (Figure S3B), we propose that the rapid differentiation of HB-EGF-deficient CD4 T cells into Th17 cells might be the most likely reason for the early onset of disease.

### CD4^ΔHB-EGF^ mice show earlier onset of EAE symptoms

To further explore whether the enhanced Th17 cell differentiation caused by HB-EGF deficiency impacts on the development of Th17 cell-driven autoimmune diseases, we induced experimental autoimmune encephalomyelitis (EAE) in C57BL/6 wt and CD4^ΔHB-EGF^ mice. In this classical mouse model of multiple sclerosis, myelin oligodendrocyte glycoprotein (MOG)-specific Th17 cells differentiate in primary and secondary lymph nodes before entering the CNS, leading to subsequent demyelination of nerves (Hirota et al., 2011). As shown in Figure 6, we found that CD4^ΔHB-EGF^ mice exhibited a significantly higher frequency of IL-17 expressing T cells following CFA/MOG_35-55_ immunization (Figure 6A) and, in comparison to their wt littermates, CD4^ΔHB-EGF^ mice exhibited an earlier onset of symptoms (Figure 6B & C).

**Figure 6:**
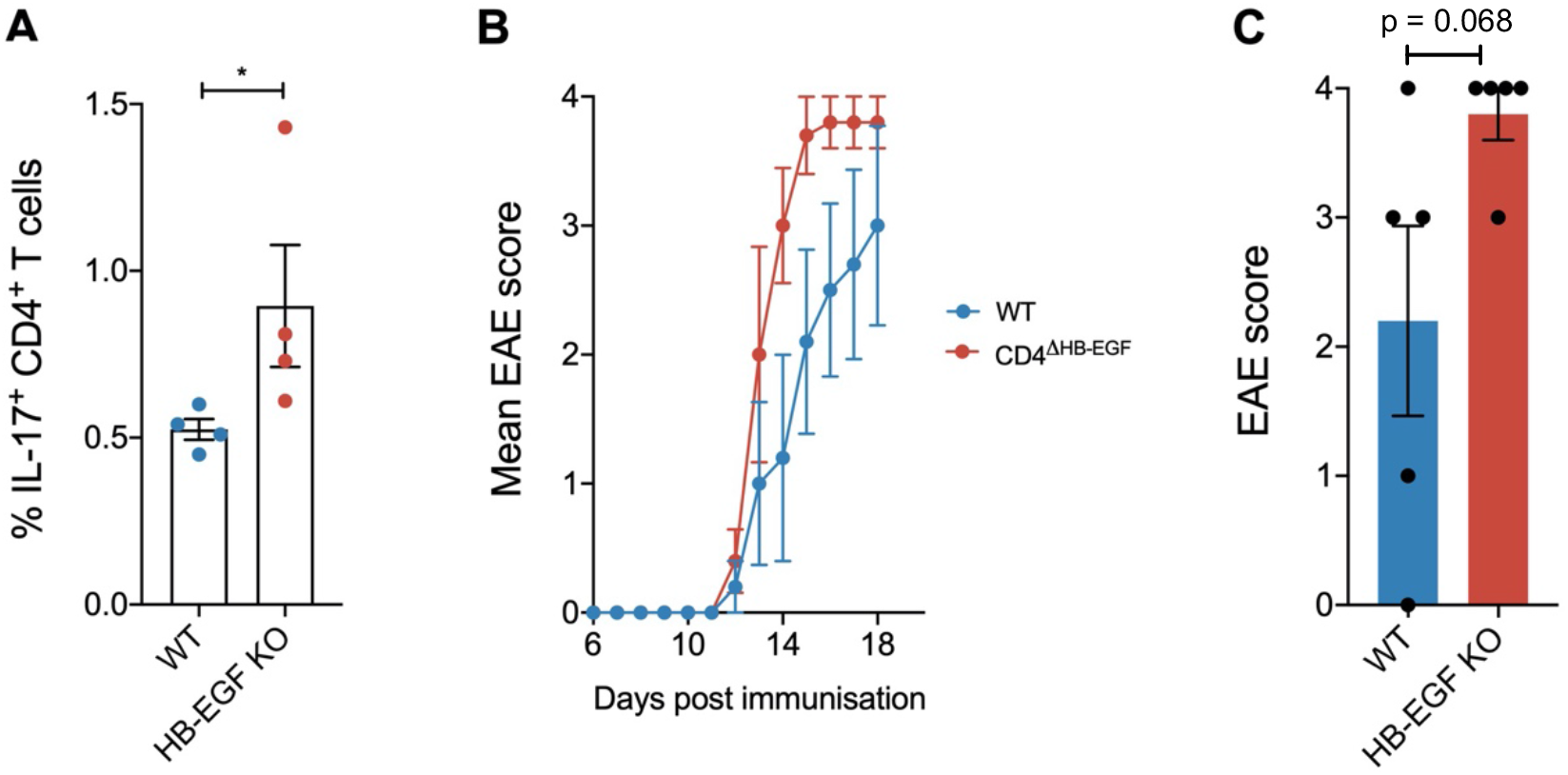
HB-EGF-deficient CD4 T cells show enhanced Th17 differentiation after immunisation with MOG_35-55_ peptide, early onset and more severe symptoms of EAE. (C) Mice were immunised with CFA/MOG_35-55_ subcutaneously into the base of the tail and culled after 7 days. Splenocytes were re-stimulated *ex vivo* with MOG_35-55_ peptide CFA/MOG_35-55_ for 6 hours. Cells were subsequently stained for CD4, CD3 plus intracellularly for IL-17. Quantified data show percentage of IL-17^+^ T cells as a proportion of CD4^+^ CD3+ lymphocytes. (B+C) EAE was induced in WT and CD4^ΔHB-EGF^ mice by subcutaneous injection of CFA/MOG_35-55_ into the base of the tail followed by intraperitoneal injection of pertussis toxin. (A) mean EAE scores and (B) EAE scores on day 15 are shown. Error bars show mean ± SD. n=4-5 in each group. Student’s t tests were used to assess significance.

Together, these findings indicate that CD4 T cell-derived HB-EGF lessens the differentiation of Th17 cells in an autocrine manner, and therefore prevents the induction of Th17-mediated inflammatory and autoimmune diseases.

## DISCUSSION

Our data reveal a novel mechanism by which T cell-derived HB-EGF constrains Th17 differentiation in an autocrine manner and thereby prevents the development of autoimmune diseases. Mechanistically, we found that T cell-derived HB-EGF induces the enhanced expression of IL-2 by CD4 T cells. IL-2 is a critical T cell survival factor. Furthermore, IL-2 induced STAT-5 signalling blocks the expression of the master Th17 transcription factor Rorγt (Laurence et al., 2007) and inhibits IL-6 receptor expression, a key cytokine signalling pathway required for the differentiation of Th17 cells (Yang et al., 2011). Nevertheless, the exact molecular mechanism by which HB-EGF diminishes Th17 differentiation remains to be addressed. It is conspicuous that of all EGFR ligands only HB-EGF has been described to induce EGFR-mediated activation of STAT-5 (Heo et al., 2018). Thus, it is tempting to speculate that HB-EGF mediated signalling may also directly sustain STAT-5 activation. In such a way, HB-EGF may in parallel to enhanced IL-2 expression also directly counterbalance IL-6 / TGFβ mediated RORγt expression and consequently Th17 differentiation. Thus, the exact contribution of the different mechanisms by which T cell-derived HB-EGF interferes with Th17 differentiation remain to be dissected in future studies.

Nonetheless, it is striking to establish a role of the EGFR in Th17 differentiation. Recently, it has been shown that the Insulin-like Growth Factor Receptor (IGFR), a transmembrane Receptor Tyrosine Kinase (RTK) closely related to the EGFR, is also expressed on activated CD4 T cells (DiToro et al., 2020). Long-term expression is retained specifically on Th17 effector cells and Tregs (DiToro et al., 2020). IGFR has been found to promote Th17 cell differentiation through activation of PLC, PKB/AKT and mTOR signalling. Conversely, our data show that HB-EGF signalling through the EGFR diminishes Th17 cell differentiation. Remarkably, the EGFR is one of the most down-regulated genes in Th17 cells differentiated in the presence of IGF (DiToro et al., 2020), suggesting that there is an antagonistic regulation of Th17 cell differentiation between the IGFR and the EGFR. As both the IGFR and EGFR are expressed on newly-activated CD4 T cells, the expression of ligands such as HB-EGF or IGF1/2 may antagonistically modulate Th17 cell differentiation.

In line with such an assumption, it has been shown previously that low strength activation of human CD4 T cells *in vitro* promotes the differentiation of cells into Th17 effector cells (Purvis et al., 2010). Our data show a direct correlation between T cell activation and HB-EGF expression. Thus, potentially a diminished HB-EGF expression might contribute to the described enhanced differentiation of low strength activated CD4 T cells into Th17 effector cells. Furthermore, in many infections, immunodominant T cell clones often express TCRs with a high affinity for their cognate antigens. Based on our data, we would assume that clones expressing low-affinity TCRs for their cognate antigen will induce a lower expression of HB-EGF than those expressing high-affinity TCRs. Such a correlation may then suggest a mechanism whereby strong TCR activation leads to sustained HB-EGF mediated IL-2 signalling and may thus favour the clonal expansion of immunodominant T cell clones. Conversely, T cell clones with a TCR showing a low affinity for self-antigen may escape central tolerance during thymic selection. Upon low-affinity antigen binding such T cells may then express minimal amounts of HB-EGF and may therefore more readily differentiate into pathogenic Th17 cells, thereby supporting the development of autoimmune diseases. Addressing such a hypothesis may require substantially more work. Nonetheless, it is striking to observe that the phenotype of HB-EGF-deficient CD4 T cells closely resembles those with a deficiency in PTPN-2 expression (Spalinger et al., 2015). Similar to HB-EGF, loss of PTPN-2 expression in T cells leads to enhanced Th17 differentiation, enhanced systemic inflammation and the transfer of naïve, PTPN-2 deficient CD4 T cells into *Rag^-/-^* mice to a rapid on-set of colitis (Spalinger et al., 2015). PTPN-2 is a pivotal regulator of EGFR activity (Stanoev et al., 2018) and several genetic studies revealed that loss-of-function variants of PTPN2 are linked to chronic inflammatory and autoimmune disorders (Doody et al., 2009). Thus, one way by which loss-of-function variants of PTPN2 may contribute to the development of autoimmune diseases could potentially be via a disrupted regulation of EGFR activity of CD4 T cells. SNPs in the promoter region of HB-EGF have been associated with Alzheimer’s Disease (AD) (Jun et al., 2017) and it has been established that AD is closely related to enhanced inflammation in the brain (Heppner et al., 2015). Therefore, it is tempting to speculate that altered HB-EGF expression by T cells might potentially be one factor contributing to AD development.

Thus, taken together, while it is already well-established that EGFR expression by CD4 T cells is a critical regulator of their effector function (Minutti et al., 2017; Zaiss et al., 2013; Zeboudj et al., 2018), our data reveal yet another pathway by which the expression of the high-affinity EGFR ligand HB-EGF prevents the differentiation of Th-17 cells. As one of the few RTKs, the EGFR shows a “biased agonism” (Freed et al., 2017). This means that ligands of different affinity induce different signals downstream of the receptor (Freed et al., 2017). The high-affinity EGFR ligand HB-EGF induces a selective autophosphorylation of EGFR-Y1068 and thus the activation of the MAPK signalling pathway (Minutti et al., 2017), which we demonstrate here leads to elevated expression levels of IL-2 and STAT-5 signalling. In contrast, the low affinity EGFR ligand Amphiregulin induces the selective autophosphorylation of EGFR-Y992 (Minutti et al., 2017) and thus a sustained signalling via the Phospholipase C (PLC) signalling pathway (Minutti et al., 2019); in this way resembling the signalling induced by the IGFR (DiToro et al., 2020). Amphiregulin-induced, sustained PLC signalling induces then the “inside out” activation of integrins, such as integrin-α_V_ (Minutti et al., 2019), a process known to mediate the local activation of bio-active TGFβ and is critical for Treg-mediated immune suppression (Worthington et al., 2015). Thus, the activation of integrin-α_V_ on Tregs might be one mechanism by which the low-affinity EGFR ligand Amphiregulin enhances suppressive capacity of Tregs (Zaiss et al., 2013). Consequently, due to the biased agonism of the EGFR the local expression of EGFR ligands with different affinity for their receptor has the potential to influence the effector function of CD4 T cells and thus may regulate local immune responses, while the interruption of this signalling pathway may predispose for the development of autoimmune diseases.

## MATERIALS AND METHODS

### Breeding of Experimental Animals

C57Bl/6, *Rag1^-/-^*, CD45.1/.2 (Ly5 hets), *CD4:cre x Hbegf^fl/fl^* and *CD4:cre x Egfr^fl/fl^* mice were bred and maintained on a C57Bl/6 background at the University of Edinburgh, under specific pathogen free conditions. Mice were 6-12 weeks of age at the start of experiments and were housed in individually-ventilated cages. Mice were both age-matched and sex-matched, with both female and male mice being used. Experiments were performed in accordance with the United Kingdom Animals (Scientific Procedures) Act of 1986. All researchers were accredited for animal handling and experimentation by the UK Home Office. Experimentation at the University of Edinburgh was approved by the University of Edinburgh Animal Welfare and Ethical Review body and granted the UK Home Office. All research was carried out under the project license PPL70/8470. No individual data points were excluded.

### Primary Cell Isolation

#### Spleen and lymph nodes

Mouse spleens and (mesenteric/inguinal) lymph nodes were homogenised by gentle processing through a 70μM cell strainer to form a single cell suspension. Splenocytes were subsequently treated with red blood cell lysis buffer (Sigma). Cells were counted using a haemocytometer and microscope (Zeiss). Staining and Cell Sorting detailed in “Flow Cytometry, Re-stimulation and FACS-sorting”.

#### Small Intestine

Small intestinal intraepithelial lymphocytes (IELs) and lamina propria lymphocytes (LPL) were isolated as described before (Minutti et al., 2017). In brief, mice were sacrificed and small intestines were placed in ice-cold complete PBS. After removal of residual mesenteric fat tissue and Peyer’s patches, intestines were cut roughly into 0.5-1cm pieces, and washed thoroughly by vortexing in PBS. Tissue was placed into pre-digestion buffer (HBSS containing 10mM HEPES, 5mM EDTA, 1mM DTT and 5% FCS) and incubated for 20 minutes at 37 degrees with frequent shaking. Tubes were vortexed intensely for 10s, and passed through 70μM cell strainers into 50ml falcon tube. The flow through contained the IEL population, and was stored on ice for further use. Tissue was transferred into fresh pre-warmed pre-digestion buffer, and incubation and straining was repeated as above. Tissues were then transferred into HBSS with 5% FCS for 10 minutes at 37 degrees with frequent shaking, vortexed, and passed through a cell strainer. All flow through containing IEL was collected and pooled together. Intestinal tissue was then transferred into pre-warmed digestion buffer (HBSS containing 10mM HEPES, 0.5mg/ml Collagenase D, 0.5mg/ml DNAse I and 5% FCS) and incubated at 37 degrees for 30 minutes with frequent shaking. Tissue was passed through a cell strainer and washed in FACS buffer (Dulbecco’s phosphate buffered saline with 2% FCS) before centrifugation at 300g for 5 minutes at 4 degrees. The cell pellet (containing LPL) was then resuspended in FACS buffer and stored on ice for further use. Staining detailed in “Flow Cytometry, Re-stimulation and FACS-sorting”.

#### Colon

Colons were removed and flushed using PBS to remove faeces. Tissue was then digested as above for small intestine. Briefly, colons were cut up into 0.5-1 cm pieces, washed by vortexing in PBS, and placed into pre-digestion buffer at 37 degrees for the specific time mentioned above. Tissues were passed through cell strainers (this time not collecting the flow through), and this process was repeated again. Tissues were incubated in HBSS with 5% FCS as above, and finally digested in digestion buffer, before centrifugation at 300g for 5 minutes at 4 degrees before resuspension of cell pellets in FACS buffer. Staining detailed in “Flow Cytometry, Re-stimulation and FACS-sorting”.

### Flow Cytometry, Re-stimulation and FACS-sorting

Single cell suspensions were prepared as previously described in “Primary Cell Isolation”. Cells were incubated with Fc block (CD16/CD32) (BioLegend), and labelled with fluorescent conjugated antibodies against different cell surface antigens (see below). For detection of intracellular markers, cells were incubated at 37 degrees for 6 hours with 2μg/ml anti-CD3 in the presence of 2μM monensin to re-stimulate cells and block protein transport, respectively. Alternatively, cells were re-stimulated by incubating with 50ng/ml phorbol myristate acetate (PMA) and 500ng/ml ionomycin at 37 degrees for 4 hours in the presence of monensin. Cells were washed and stained for surface markers as above, before subsequent fixation with 2% paraformaldehyde in Dulbecco’s phosphate buffered saline for 20 minutes at room temperature, permeabilisation with Perm wash (BD Biosciences), and staining with antibodies against intracellular antigens. For detection of intranuclear antigens, cells were stained for surface markers, fixed and permeabilised using FoxP3 staining buffer set (eBioscience), and subsequently stained with antibodies against intranuclear antigens. For Flow Cytometry, Live/Dead (Life Technologies) was used to exclude dead cells from analysis, and samples were analysed using Becton-Dickinson BD SORP LSR II and FlowJo software v9 (Treestar Inc). For Cell Sorting, naïve cells (CD4^+^ CD25^-^ CD45RB^hi^) were sorted using BD FACSAria IIU and again analysed using FlowJo software.

### In vitro T cell differentiation

Sorted naïve CD4 T cells (see “Flow Cytometry and FACS”) were cultured in Iscoves Modified Dulbecco’s Medium (IMDM) containing 10% FCS, penicillin/streptomycin, L-Glutamine and β-mercaptoethanol (from now referred to as ‘complete IMDM’) at a density of 10^6^/ml. For Th1 and iTreg differentiation, cells were cultured in plates coated with 1 μg/ml anti-CD3 and 2μg/ml anti-CD28. For Th17 differentiation, cells were cultured in plates coated with 1μg/ml anti-CD3 and 10μg/ml anti-CD28. The following cytokines were added to generate each subset: **Th1** 10ng/ml IL-12 and 5μg/ml anti-IL-4 (clone 11B11); **iTreg** 10ng/ml TGFβ and 10ng/ml IL-2; **Th17** 50ng/ml IL-6, 10ng/ml IL-1β, 10ng/ml IL-23, 1ng/ml TGFβ and 300nM FICZ. After 48 hours, 5ng/ml IL-2 was added to all wells for Th1 differentiation. Recombinant HB-EGF was added to cultures as specified (0.01–100ng/ml). Cells were split 1:2 on day 2 after stimulation and analysed by Flow Cytometry on day 4-6.

### Cell Trace Violet proliferation

Sorted naïve CD4 T cells were stained with 1μl/ml CellTrace Violet Proliferation Kit (Thermofisher, Cat. C34571) in the dark for 20 minutes at room temperature. Five times the volume of complete IMDM was subsequently added for 5 minutes at room temperature before centrifugation. Cells were cultured at a density of 10^6^/ml in pre-warmed complete IMDM containing mouse T-activator anti-CD3/CD28 Dynabeads (Thermofisher) to give a final bead-to-cell ratio of 1:1. After 24 hours, some cells were analysed for CellTrace Violet expression to generate a baseline peak. After 48 hours, 2ng/ml IL-2 was added to cells. After 96 hours, all cells were analysed for CellTrace Violet expression.

### IL-2 ELISA

Sorted naïve CD4 T cells were cultured in complete IMDM at a density of 10^6^/ml in the presence of mouse T-activator anti-CD3/CD28 Dynabeads (Thermofisher) at a bead-to-cell ratio of 1:1. After 24 hours, cells were centrifuged and supernatants stored at −20 degrees for further analysis by ELISA. IL-2 ELISA was performed using IL-2 Mouse Uncoated ELISA Kit (Invitrogen) following manufacturer’s protocol.

### Anti-CD3 injection

Mice were injected intraperitoneally with 50μg anti-CD3 antibody (clone 145-2C11) in PBS. After 48 hours, mice were sacrificed and spleens, mesenteric lymph nodes and small intestines harvested. Cells were isolated as specified in “Primary Cell Isolation” and re-stimulated/stained as detailed in “Flow Cytometry, Re-stimulation and FACS-sorting”.

### *Rag1^-/-^* T cell transfer model and TGFβ blocking antibody addition

*Rag1^-/-^* CD45.1 mice were injected intravenously with 4×10^5^ FACS sorted naïve CD4^+^ T cells. Depending on the experiment, mice were either injected with wild type or HB-EGF-deficient cells, or a 1:1 mixture of wild type (CD45.1) and HB-EGF-deficient (CD45.2) cells. In specified experiments, *Rag1^-/-^* mice were injected with 5mg/kg TGFβ blocking antibody (1D11) 3 times weekly for the first 3 weeks. Wasting disease was determined by weight loss compared to original weight, and mice culled when the group lost an average of 20% body weight. To assess cell proliferation and effector CD4^+^ T cell populations in *Rag1^-/-^* mice receiving a 1:1 mix of cells, mice were culled at weeks 1, 2 and 3 post injection of naïve CD4^+^ T cells. Cells were isolated and stained from spleens, mesenteric lymph nodes and colons as specified in “Primary Cell Isolation” and “Flow Cytometry, Re-stimulation and FACS-sorting”. Spleens, liver, kidney, pancreas, small intestines and colons were placed into 10% neutral-buffered Formalin solution overnight, before transfer into 70% ethanol solution for analysis by histology. Colons and small intestines were stained with heamatoxylin and eosin (H&E).

### EAE

Mice were anaesthetised using Isoflurane and immunised subcutaneously with 200μg MOG_35-55_ (MEVGWYRSPFSRVVHLYRNGK) emulsified in Complete Freund’s Adjuvant (CFA) containing 500μg/ml heat-killed Mycobacterium tuberculosis (H37RA), into both hind legs. Mice received 200ng pertussis toxin intraperitoneally on days 0 and 2 after induction. Clinical evaluation was performed using a 5-point scale: **0** no clinical signs; **1** flaccid tail; **2** impaired righting reflex and/or gait; **3** partial hind limb paralysis; **4** total hind limb paralysis; **5** total hind limb paralysis with partial fore limb paralysis. Mice were culled once they exceeded grade 3, or upon >20% weight loss or adverse signs such as substantial dyspnoea, weakness, dehydration or a hunched appearance.

### Human CRISPR

Human CD4 T cells were MACS-sorted from PBMC and subsequently pre-activated with α-CD3/CD28 Dynabeads (Thermofisher) at a bead-to-cell ratio of 1:1 for 48 hours. Next, dynabeads were removed and the CD4 T cells were rested for 24 hours. For CRISPR/Cas9 knockout 0.8×10^6^ cells were mixed with 2.5 μl crRNA/TracRNA complex (200uM), 1.67 μl Cas9 nuclease 3NLs and 5.83 μl Buffer R (integrated DNA technologies, Coralville, US) and incubated at room temperature for 20 min. This cell mix was transfected using the neon pipette station 1600 volts, 10 ms, 3 pulses. After electroporation cells were transferred into 48-wells plate containing pre-warmed complete RPMI culture media and cultured for 24 hours before analysis of Th17 cell differentiation. Sequences for crRNA guides were as follows: *Hbegf* target sequences ACTGGCCACACCAAACAAGGAGG, GACCAGCTGCTACCCCTAGGAGG and CTTCATGGTCCCGCACCGAGAGG; *Egfr* target sequences GCTGCCCCGGCCGTCCCGGAGGG, GAGGATGTTCAATAACTGTGAGG and GAAAACCTGCAGATCATCAGAGG.

### Statistical analysis

Statistical analysis was performed using Prism (GraphPad Software). All data were analysed using the Student’s *t* test. P-values <0.05 were considered statistically significant.

## ACKNOWLEDGEMENTS

We thank N. Logan and the other members of the Zamoyska Lab for all their help and excellent technical assistance; and the vivarium support staff for excellent animal husbandry. D.M.Z. was supported by the Medical Research Council, grant MR/M011755/1. R.Z. has been supported by a Wellcome Trust Investigator Award (WT205014/Z/16/Z) and F.M. by an EastBio BBSRC PhD fellowship.

## AUTHOR CONTRIBUTIONS

F.M. designed and performed experiments, analysed and interpreted data, and wrote the manuscript. K.P., A.M., R.C.S, L.W.P., L.N., J.Y. and D.H. performed experiments. D.W. provided expertise in CRISPR/Cas9 experimental design. S.H. produced the TGFβ blocking antibodies. A.G. stained sections and provided expertise in histology. M.W. provided expertise in Flow Cytometry and FACS, and edited the manuscript. R.Z. and J.V.L. contributed tools, provided expertise, and edited the manuscript. D.M.Z. designed the research, interpreted data and wrote the manuscript.

## Notes

### Competing Interest Statement

The authors have declared no competing interest.

