## Supplementary Figures for "T cell derived HB-EGF prevents Th17 cell differentiation in an autocrine way"

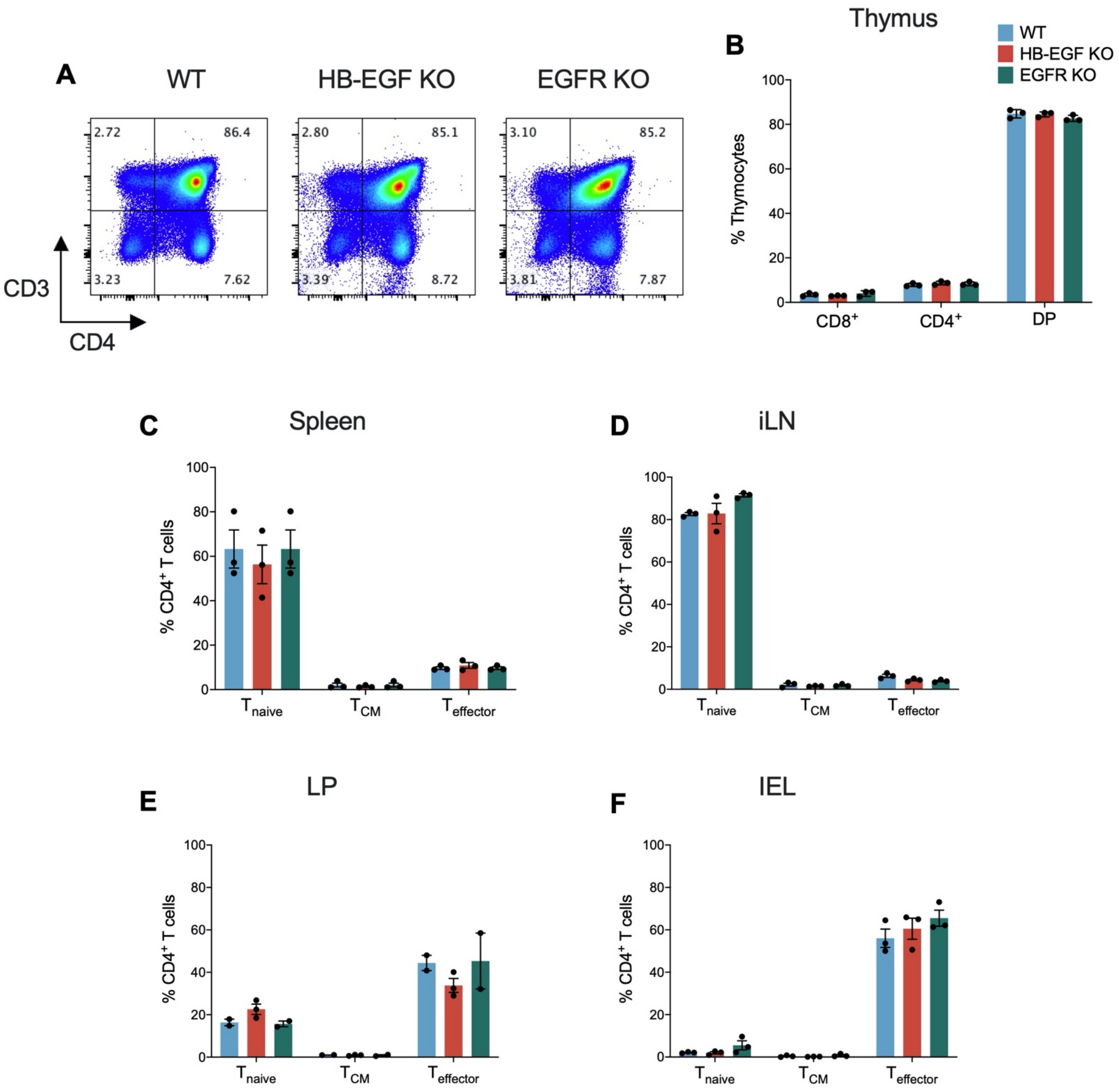


**Supplementary Figure 1: Steady state characterisation of CD4$^{\Delta HB\text{-}EGF}$ mice.** Surface staining (A) and quantification (B) for CD4/CD8/double positive (DP) cells from thymus of WT, CD4$^{\Delta HB\text{-}EGF}$ (referred to as HB-EGF KO) and CD4$^{\Delta EGFR}$ (referred to as EGFR KO) mice. Numbers in quadrants indicate percentage of cells stained as a total of thymocytes.

Splenocytes (C), Inguinal Lymph Nodes (iLN) (D), Lamina Propria (LP) lymphocytes (E) and Intraepithelial (IEL) lymphocytes (F) from WT, CD4$^{\Delta HB\text{-}EGF}$ and CD4$^{\Delta EGFR}$ mice were stained for CD4, CD3, CD62L and CD44 and analysed by flow cytometry. Quantified graphs show percentages of naïve (CD62L$^{+}$ CD44$^{-}$), central memory (CD62L$^{+}$ CD44$^{+}$) and effector (CD62L$^{-}$ CD44$^{+}$) T cells as a proportion of CD3$^{+}$ CD4$^{+}$ lymphocytes from the respective organs. Error bars show mean ± SEM. n=3 in each group.

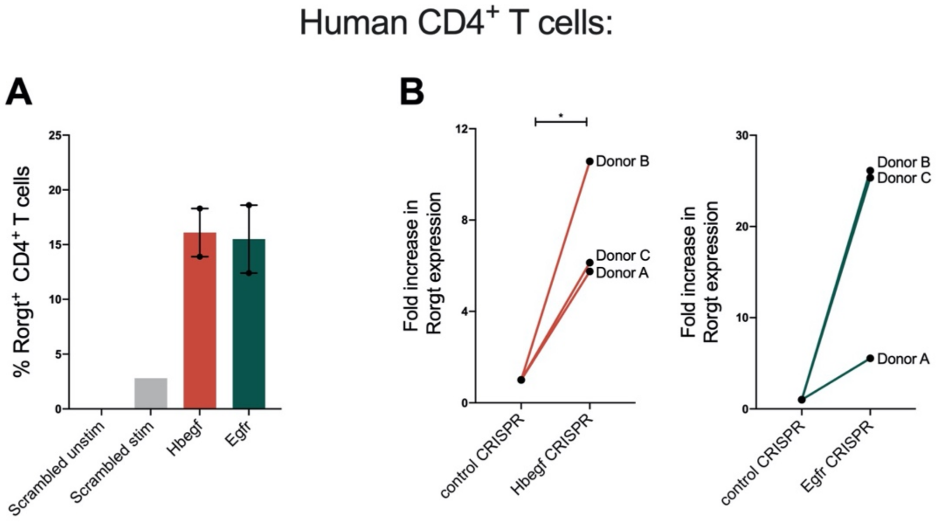


**Supplementary Figure 2: Enhanced *in vitro* Th17 cell differentiation of HB-EGF-deficient CD4+ T cells in humans.** $CD4^+$ T cells were MACS sorted from PBMCs isolated from healthy donor human blood. Cells were transfected with CRISPR/Cas9 guide RNA specific for *Hbegf* and *Egfr* genes in order to delete these genes, or a scrambled guide RNA sequence used as a negative control. Cells were rested in recombinant IL-2 for 2 days, before activation and polarisation towards a Th17 cell phenotype. (E) Quantification of Ror$\gamma$t$^+$ cells as a proportion of $CD4^+$ lymphocytes from unstimulated cells, cells transfected with scrambled guide RNA (control), or cells transfected with *Hbegf* or *Egfr* guide RNA from Donor A. (F) Fold increase in Ror$\gamma$t expression from cells transfected with *Hbegf* (left) or *Egfr* (right) guide RNA compared to cells transfected with scrambled guide RNA for Donors A-C. Each experiment was performed at least twice with similar results. Error bars show mean $\pm$ SEM. n=2-3 in each group. Student's t tests were used to assess significance.

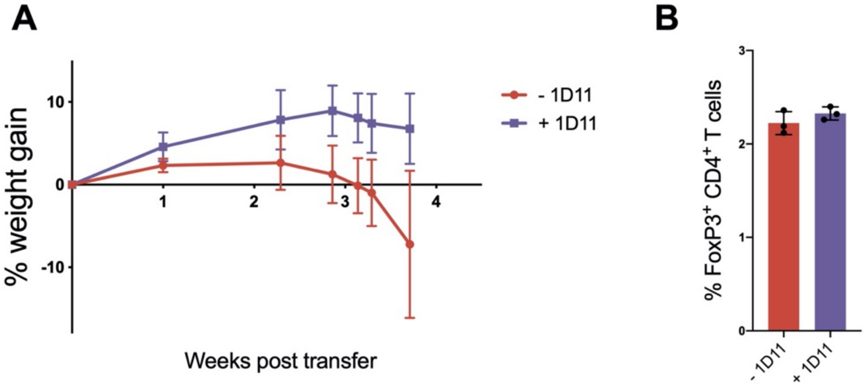


**Supplementary Figure 3: TGFβ blockade prevents early onset of colitis.** (E+F) *Rag1*$^{-/-}$ mice were intravenously injected with 500,000 HB-EGF-deficient naïve CD4 T cells and administered TGFβ blocking antibody 1D11 (purple) three times a week for three weeks or left untreated (red). (E) Graph shows percentage weight gain of mice compared to their original body weight post injection. (F) Upon onset of wasting disease mice were culled and spleens stained for CD4, CD3 and FoxP3. Quantified data shows percentage FoxP3$^{+}$ T cells as a proportion of CD4$^{+}$ lymphocytes. Error bars show mean ± SEM. n=3-5 in each group. Student's t tests were used to assess significance.
